# Neural circuit basis of adolescent THC-induced potentiation of opioid responses later in life

**DOI:** 10.1101/2024.04.23.590773

**Authors:** Elizabeth Hubbard, Vivienne Mae Galinato, Pieter Derdeyn, Katrina Bartas, Stephen V. Mahler, Kevin T. Beier

**Affiliations:** Department of Physiology and Biophysics, University of California, Irvine, Irvine, CA, USA 92697-4560; Program in Mathematical, Computational, and Systems Biology. University of California, Irvine, Irvine, CA, USA 92697-4560; Department of Neurobiology and Behavior, University of California, Irvine, Irvine, CA, USA 92697-4560; Department of Biomedical Engineering, University of California, Irvine, Irvine, CA, USA 92697-4560; Department of Pharmaceutical Sciences, University of California, Irvine, Irvine, CA, USA 92697-4560

## Abstract

Use of one drug of abuse typically influences the behavioral response to other drugs, either administered at the same time or a subsequent time point. The nature of the drugs being used, as well as the timing and dosing, also influence how these drugs interact. Here, we tested the effects of adolescent THC exposure on the development of morphine-induced behavioral adaptations following repeated morphine exposure during adulthood. We found that adolescent THC administration potentiated reward, and paradoxically impaired the development of morphine reward. Following a forced abstinence period, we then mapped the whole-brain response to a moderate dose of morphine, and found that adolescent THC administration led to increased morphine-induced activity in the basal ganglia and increased functional connectivity between frontal cortical regions and the ventral tegmental area. Last, we show using rabies virus-based circuit mapping that adolescent THC exposure triggers a long-lasting elevation in connectivity from the frontal cortex regions onto ventral tegmental dopamine cells. Our study adds to the rich literature on the interaction between drugs of abuse, including THC and opioids, and provides potential circuit substates by which adolescent THC exposure influences responses to morphine later in life.

## INTRODUCTION

Cannabis is currently the most widely used drug by adolescents in the United States^1^. Despite a reputation as a relatively harmless drug, over 30% of cannabis users meet the diagnostic criteria for cannabis use disorder (CUD)^2^, a similar proportion to those who become addicted after trying “hard” drugs such as cocaine and opioids^3^. With the global trend toward cannabis legalization enabling increased access to cannabis, it is critical to understand how adolescent cannabis use impacts the brain and behavior, especially if it does so in a persistent manner in the dynamically developing adolescent brain. Adolescent exposure to the major psychoactive constituent of cannabis, tetrahydrocannabinol (THC), has been linked to a variety of negative outcomes later in life^4–6^, including increased subsequent consumption of ‘harder” drugs^7–12^. Given the posited role of the mesolimbic dopamine system as the final common pathway for addiction^13^, it is not surprising that brain regions such as the ventral tegmental area (VTA), medial prefrontal cortex (mPFC), and nucleus accumbens (NAc) are critically important in mediating the long-lasting effects of adolescent THC on responses to psychoactive drugs later in life^10,14–16^. Much like use of other psychoactive drugs^17–20^, adolescent THC usage can lead to a hyperdopaminergic state^15^ that then facilitates a variety of neuroadaptations that promote addiction. Despite this common framework, our mechanistic understanding of how adolescent THC causes persistent changes that linger until adulthood is limited.

Given that THC carries an addictive liability, it is critically important to understand the interactions between adolescent THC usage and exposure to drugs of abuse later in life, for example prescription opioids. It has long been hypothesized that cannabis use may predispose individuals to opioid use^21–23^. This is supported by the observation that cannabinoids and opioids may both activate the midbrain dopamine system, perhaps via common mechanisms of mu opioid receptor activation^24^. In addition, cross-sensitization of THC and morphine has been demonstrated^25^, adolescent THC exposure can alter expression of genes related to the endogenous opioid system^10^, as well as potentiate opioid-seeking behaviors later in life^26,27^, providing further evidence that these drugs may work via common mechanisms. This is an important question because, despite the goal to reduce the clinical usage of opioids, opioids remain the primary treatment for post-operative pain management. In addition, approximately eighty percent of those who use heroin began using opioids for pain management^28,29^. Therefore, it is important to understanding mitigate the negative effects that adolescent THC may have on behavioral responses to opioids later in life.

Here, we administer a moderate dose of THC (5 mg/kg) for 14 days starting at postnatal day 30 (PD30), during adolescence. After a washout period during which mice develop to adulthood, we then test a variety of opioid-induced behaviors by administering morphine and assessing differential effects in those with or without a history of adolescent THC. After undergoing a forced abstinence period following repeated morphine injections, we also assess how adolescent THC administration influences brain-wide activity patterns following a moderate of morphine, as well as how adolescent THC administration alters connectivity onto dopamine neurons in the VTA.

## METHODS

All procedures were approved by the University of California, Irvine’s Institutional Animal Care and Use Committee (IACUC) and carried out in accordance with the National Institutes of Health guidelines for the care and use of animals. Adolescent male and female C57BL/6J mice were weaned and sexed at PD21 and group housed with same-sex littermates. Mice were housed in standard individually ventilated cages with corncob bedding and two cotton nestlets for enrichment. Lights were on a 12 hr on/12 hr off cycle (7:30-7:30) and rooms have controlled temperature (22°C +/- 2°C) and humidity (55-65%). Mice received ad libitum access to food and water. Mice were housed in the Gillespie Neuroscience Research Facility (GNRF), except for during adolescent THC treatment. Mice were transferred to the McGaugh Hall mouse facility following weaning at approximately P25, and then returned to the GNRF at approximately P45.

### Adolescent THC Administration

Mice were handled and weighed daily. On PD30, mice began daily injections of either 5 mg/kg THC or vehicle, which was prepared daily by dissolving in 5% Tween80 in saline. After 14 days of daily injections (PD30-PD43), mice were left in their home cage undisturbed (aside from weekly cage changes) until PD70 when behavioral testing began.

### Drugs

THC was provided by the NIDA Drug Supply Program or Cayman Chemicals (Ann Arbor, MI) and administered at 5 mg/kg. Morphine was purchased from Patterson and was administered at 10 mg/kg.

### Behavioral Testing

### Locomotion

The distance traveled in the open field was assessed during 30-minute recording sessions. Locomotion was assessed once per day for seven days, the first two following saline injections for habituation days, and the last five following injection of 10 mg/kg morphine.

### Conditioned place preference

Custom two chamber acrylic boxes separated by a small corridor with distinct visual contexts on the walls of each side were used for CPP tests. On the pre-test day, mice were placed in the box with no barriers and were allowed to freely explore both sides for 30 minutes, and the time spent in each chamber was recorded. Following the pre-test, mice underwent two saline pairings and two morphine pairings. In the morning, the mice were given saline and paired with the preferred chamber, and in the afternoon, injected with morphine and paired with the non-preferred chamber. Control mice received saline in both contexts. On the post-test day, mice were again placed in the CPP chamber with no barriers, and the amount of time spent in each chamber was recorded. Data were plotted as the relative time spent on the drug-paired chamber in the post-test relative to the pre-test. Mice that spent over 1200 seconds on one side during the pre-test were excluded from analysis.

### Elevated plus maze

Mice were placed at the end of one of the open arms, facing outwards. Mice were then able to freely explore the maze for 5 minutes, and the time spent in the open arms was recorded. Tests were performed in dim lighting ∼25 lux.

### Open field test

Mice were placed in a 40cm × 40cm square arena made of opaque plexiglass and were able to freely explore during a 5-minute test. During analysis, the arena was divided into nine equal square zones. The time spent in the center square for each mouse was calculated. Tests were performed under bright lighting conditions of ∼1000 lux.

### Marble burying

Mice were placed in a rat cage with 20 marbles placed on top of corn-cob bedding. After 30 minutes, the number of marbles that were buried by each mouse was recorded.

### Von Frey

The von Frey test was used to assess mechanical sensitivity. Mice were placed on top of a wire mesh platform, restrained underneath a glass beaker, and allowed to acclimate for at least 30 minutes. The mouse’s right hindpaw was gently touched with different filaments, starting with the 0.4g filament. If the mouse showed a withdrawal response (such as lifting, shaking, or licking the paw), the experimenter stepped down to a lighter filament, and if the mouse did not respond, the experimenter stepped up to a thicker filament until a response occurred. The process continued with filaments of higher or lower forces for four additional turns following the first change in response. This up/down method was used to determine the mechanical force required to elicit a paw withdrawal response (50% threshold)^30^.

### cFos induction and immunohistochemistry

One hour following injection of 5 mg/kg morphine (a dose used only for morphine-induced reinstatement), mice were perfused with PBS followed by 4% formaldehyde, and brains isolated and post-fixed overnight in 4% formaldehyde. The following day, brain clearing began using iDISCO^31^. The monoclonal rat anti-cFos antibody (Synaptic Systems lot #226017) in 5% NDS/PBS-Tween 0.1% was used at a 1:5000 dilution, with an incubation time of five days. Donkey anti-rat Cy3 (Jackson Immunoresearch) was then used at a dilution of 1:250. Brains were then cleared in dibenzyl ether and imaged using a Zeiss Z1 light-sheet microscope. ClearMap^32^ was then used to align the imaged brains to the Allen Brain Reference Atlas, and to quantify the number of cFos+ neurons in anatomically-defined brain regions.

### Network Analysis

In order to relate functional connectivity using cFos labeling to anatomical connectivity, anatomical projection data from the Allen Mouse Brain Connectivity Atlas was used to construct a connectivity matrix among the brain regions in our analysis^33^. This atlas quantifies axonal arborization from AAV injections to measure projections from one region to another. The density of each projection, or the sum or projecting pixels divided by the total pixels in that region, was normalized by dividing by the volume of injection, so as not to bias experiments with more virus injection than others. This quantity was also multiplied by the number of voxels in the injection structure to avoid overrepresentation of smaller regions. This quantity was computed for every pair of source and target regions to build a connectivity matrix. We log-normalized these quantities so that they would have a more normal distribution and behave better in network analysis. Community detection was performed using the Louvain algorithm with a resolution parameter of 1, in the scit-kit network library^34^. Once the clusters were determined, the cFos fold change of their regions were compared using a one-way ANOVA followed by pairwise t-tests.

### Functional Analysis

Regional cFos counts across each mouse in the adolescent vehicle-treated and THC-treated conditions were dimensionally reduced using UMAP, so that regions with similar counts across the mice would be embedded close to each other^35^. Regions were clustered using HDBSCAN, which can identify hierarchical clusters in continuous data based on variations in density of the data^36^. Hierarchical clustering was also performed on the correlations between regions, to look more closely at their functional relationships. R^2^ value from Pearson correlations were computed for each pair of regions, based on their counts across mice of a given condition. Hierarchical clusterings were computed on correlation matrices using the average linkages methodology. For visualizations, the hierarchical clustering tree was cut at a height of 0.75 to determine the clusters (Figure 4I-J). The trees were cut for heights between 0.5 and 1 to observe trends in modularity across the two conditions. The seaborn library was used for all hierarchical clustering computations^37^.

### Rabies virus circuit mapping

Rabies virus (RABV) monosynaptic tracing studies were carried out as previously described^19,38^, with minor modifications. On Day 0, 100 nL of a 1:1 volume mixture of AAV-CAG-FLEx^loxP^-TC and AAV-CAG-FLEx^loxP^-G was injected into the VTA of PD56 mice that had undergone two weeks of THC injections between PD30 and PD43. Fourteen days later, RABV was injected into the VTA. After recovery, mice were housed in a BSL2 facility for 5 days to allow RABV spread and GFP expression. Mice were then perfused with PBS followed by 4% formaldehyde and postfixed overnight. Fixed brains were cryoprotected by incubation in a 30% sucrose solution overnight, and frozen in OCT. Brains were then serially sliced on a cryostat into 60 um sections. GFP+ cells throughout the brain in 22 anatomically-defined input sites were counted manually, as previously performed^19,38^.

### Viruses and injection coordinates

Coordinates used for the VTA were (relative to Bregma, midline, or dorsal brain surface and in mm): AP − 3.20, ML 0.4, DV − 4.2.

The titers of viruses, based on quantitative PCR analysis, were as follows: AAV-CAG-FLEx^loxP^-TC, serotype 5, 2.4 × 10^13^ genome copies (gc)/ml

AAV-CAG-FLEx^loxP^-G, serotype 8, 1.0 × 10^12^ gc/ml

For RABV, the titer was performed on HEK 293T-TVA800 cells.

RABV (EnvA-pseudotyped, GFP-expressing), 5.0 × 10^8^ colony forming units (cfu)/ml

After scaling RABV data, PCA was applied so features were brain regions and observations were individual brains of mice treated with various drugs. Differences between mouse brains were visualized by plotting PC1, PC2, and PC3 (Figure 5J-L, Q-R). Ellipsoids for each condition were calculated and overlayed onto plots to visualize trends in PC values by condition. To further quantify relationships between brains and conditions, the Euclidean distance between each point (or each brain) in the PC1 versus PC2 coordinate space was calculated. These distances were then plotted in a heatmap with brains being organized by similarity assessed by hierarchical clustering (Figure 5M). Alternatively, for a summary by condition, these distances were averaged based on drug administered, and the resulting heatmap and clustering were made based on averaged Euclidean distance values (Figure 5S). Heatmaps and clustering were made using the clustermap function in the Python package seaborn with default parameters^37^. The RABV data that included single injections of psychostimulants, morphine, or nicotine, or two anesthetic doses of ketamine/xylazine, were presented in our published work^39^ and used here for sake of comparison. For these experiments, a single dose of 15 mg/kg cocaine, 1 mg/kg amphetamine, 2 mg/kg methamphetamine, 0.5 mg/kg nicotine, or 10 mg/kg morphine were given to adult mice, and RABV tracing was performed the following day. These mice were anesthetized with isoflurane-based anesthesia, as used in this manuscript. A separate group was anesthetized with a ketamine/xylazine mixture prior to both helper virus injection and RABV injection, and not given another drug. RABV was introduced during the second ketamine/xylazine injection.

### Quantification and Statistical Analysis

Statistics for all studies were calculated using GraphPad Prism 9 software. Statistical significance between direct comparisons was assessed by unpaired or paired t-tests. When multiple conditions were compared, one- or two-way ANOVAs were first performed, as appropriate, then t-tests were then performed for each individual comparison. Multiple comparisons corrections were performed when multiple such t-tests were being performed e.g., RABV input mapping and cFos quantification, and significance was assessed using the Holm-Bonferroni method. In conditions where multiple comparisons were performed and the results were still considered significant, asterisks were presented corresponding to the original p-values. Paired t-tests were used in cases with repeat measurements. Dot plots presented throughout the manuscript include a bar representing the mean value for each group. Error bars represent s.e.m. throughout. For all figures, ns P > 0.05, * P ≤ 0.05, ** P ≤ 0.01, *** P ≤ 0.001, **** P ≤ 0.0001.

## RESULTS

### Adolescent THC exposure history reduces morphine-induced locomotion, with no impact on morphine reward

We first exposed mice to a daily 5 mg/kg/day dose of THC (or its vehicle) during adolescence, here defined as postnatal days 30-43 (Figure 1A). This is considered a moderate dose that is comparable to doses self-administered by humans^40,41^. Following vehicle or THC administration, mice were left in their home cages undisturbed besides weekly cage changes until postnatal day 70, at which time behavioral testing commenced. The protocol of morphine administration and behavioral testing is shown in Figure 1B. To test how adolescent THC exposure history impacts morphine reward, we conducted a conditioned place preference (CPP) test using 10 mg/kg morphine (Figure 1C). We induced a significant preference to the morphine-paired side in both vehicle and THC exposed animals, to an approximately equivalent extent. No preference was observed in animals treated with vehicle during adolescence and again at adulthood in place of morphine (Vehicle/saline 762s pre, 778s post, p = 0.22; Vehicle/morphine 783s pre vs. 887s post, p = 0.038; THC/morphine 785s pre vs. 887s post, p = 0.0017; Figure 1D). To determine whether adolescent THC treatment impacted morphine-induced locomotion, we measured locomotor behavior while mice were in an open field. Following a two-day habituation protocol where mice were given saline injections, 10 mg/kg morphine was given once per day for five days (Figure 1E) to test the sensitization or tolerance to morphine’s effects on locomotion. We observed that morphine increased locomotion the most following the first dose, and this elevation was reduced with each subsequent dose, for both groups. We also found that that adolescent THC exposure overall blunted morphine-induced locomotion during the five-day morphine administration window (repeated measures Two-way ANOVA p = 0.045; Figure 1F), though no individual day was significantly different between the groups following multiple comparisons corrections. These data indicate that adolescent THC exposure reduced morphine-induced locomotion following morphine exposure later in life.

**Figure 1:**
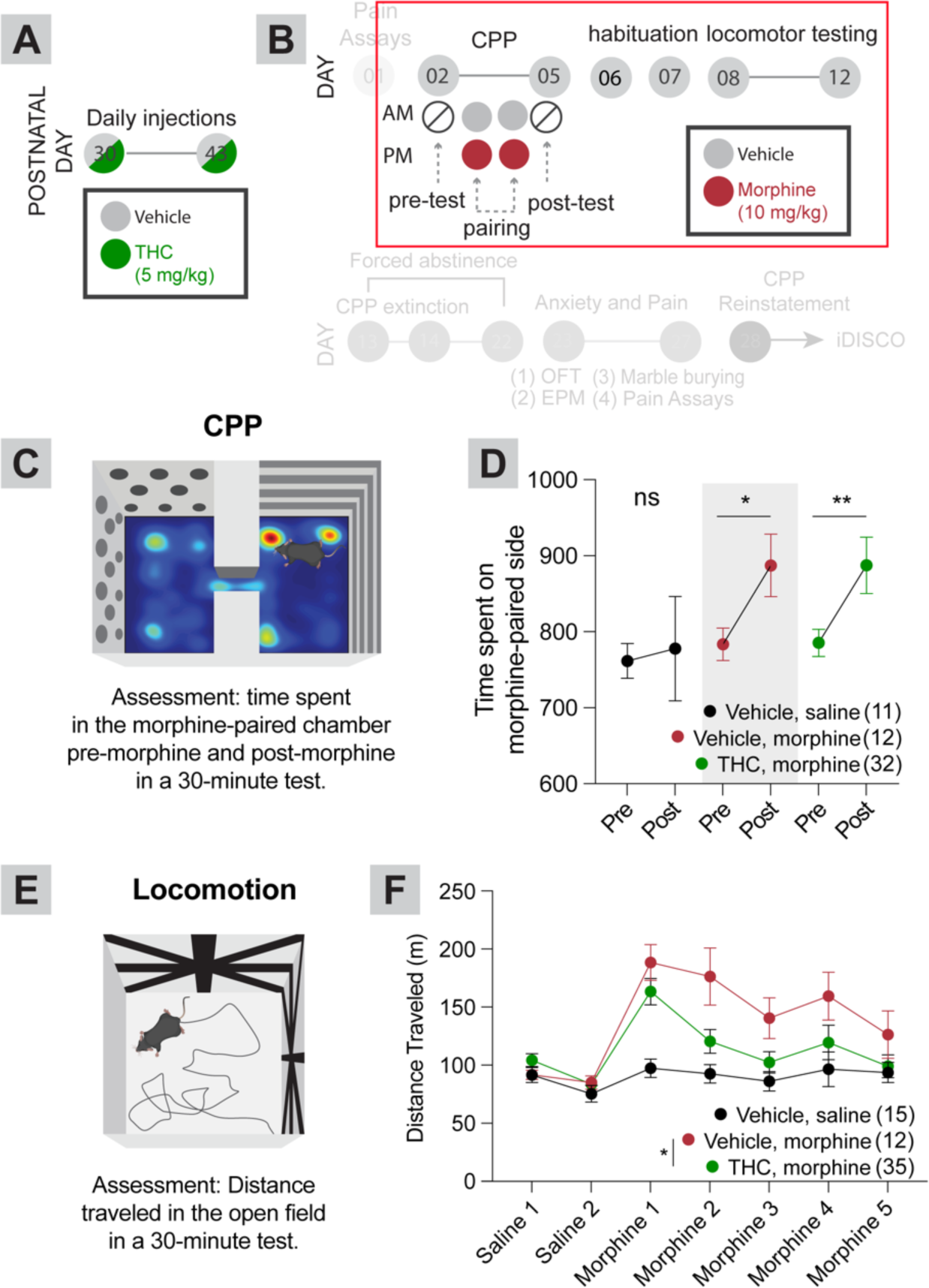
Effects of adolescent THC exposure on behavioral responses to morphine administration later in life. **A,** Timeline of THC or vehicle injections during adolescence. Injections were repeated once per day for 14 total days. **B**, Morphine injections, and behavioral testing. The entire protocol is shown for each of Figures 1-3 for reference, with the relevant tests presented in the figure in full opacity and surrounded by red boxes. **C**, Schematic of the conditioned place preference test. **D**, Time spent in the morphine-paired chamber in the pre-test and post-test (in the vehicle/saline-paired group, both chambers were paired with saline). Vehicle/saline 762s pre, 778s post, p = 0.22; Vehicle/morphine 783s pre vs. 887s post, p = 0.038; THC/morphine 785s pre vs. 887s post, p = 0.0017. **E**, Schematic of the locomotor test. **F**, Mice treated with THC during adolescence showed an overall reduction in morphine-induced locomotion relative to mice treated with saline during adolescence. Repeat measures Two-way ANOVA between conditions p = 0.045; Sidak’s multiple comparisons tests morphine day 1 p = 0.68, morphine day 2 p = 0.24, morphine day 3 p = 0.31, morphine day 4 p = 0.50, morphine day 5 p = 0.75. Vehicle/saline-treated mice are shown for comparison but were not included in statistical tests. Two-way ANOVA only included morphine injection days, not habituation days. For this and all figures, error bars = SEM, ns p > 0.05, * p < 0.05, ** p < 0.01, *** p < 0.001, **** p < 0.0001.

**Figure 2:**
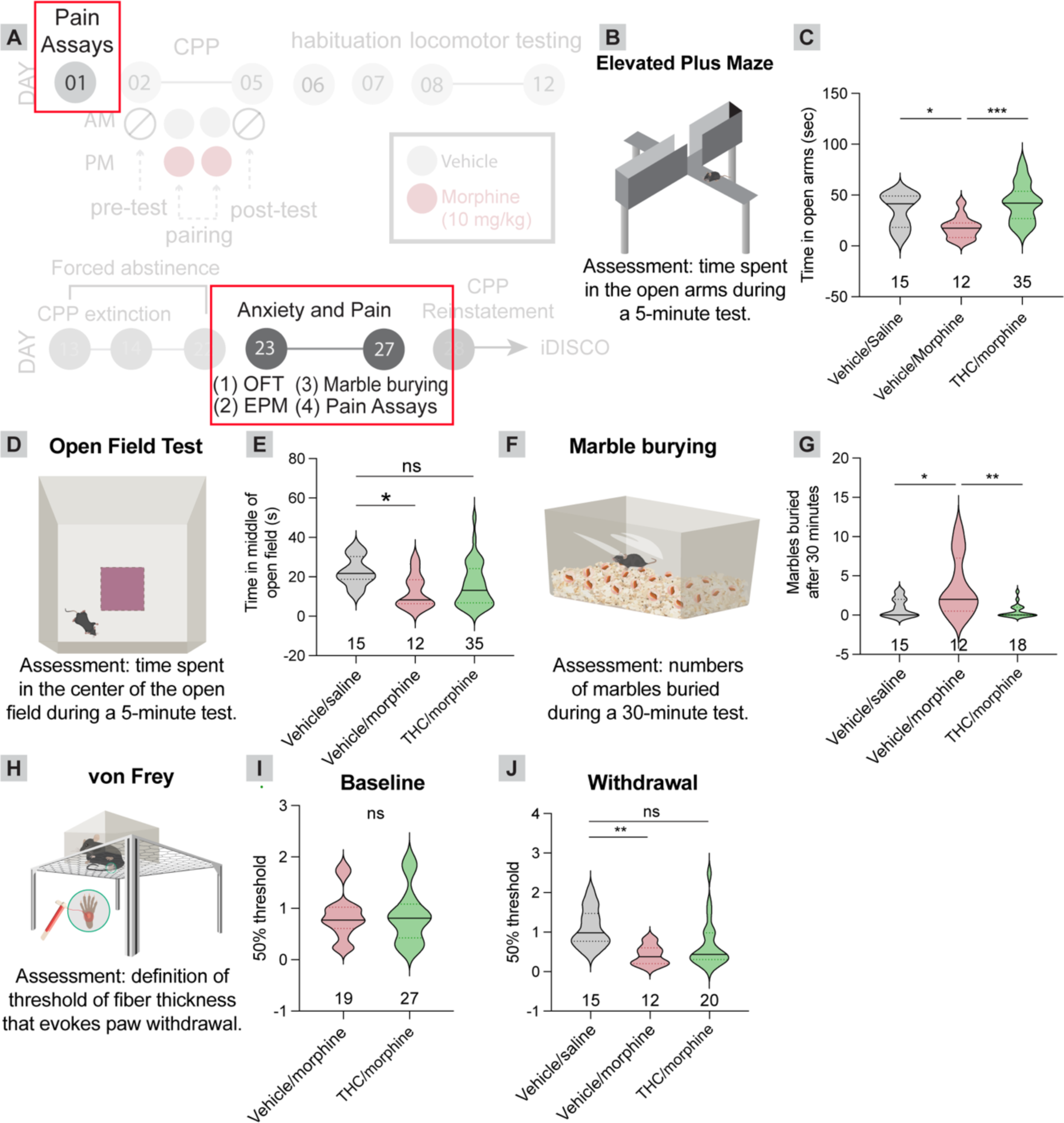
Adolescent THC administration reduces expression of morphine-induced withdrawal behaviors during a prolonged abstinence period. **A**, Timeline of experiments. **B**, Schematic of the elevated plus maze test. **C**, Repeated morphine administration leads to a reduction in the time spent in the open arms of an elevated plus maze, indicative of anxiety-like behavior relative to mice treated with saline during adolescence, and saline during adulthood. Adolescent THC exposure prevented this morphine-induced anxiety. One-way ANOVA p = 0.0005; pairwise t-tests, vehicle/saline vs. vehicle/morphine 47.73 s vs. 17.84 s, p = 0.01; vehicle/saline vs. THC/morphine 47.73 s vs. 42.39 s, p = 0.67; vehicle/morphine vs. THC/morphine 17.84 s vs. 42.39 s, p = 0.0018. **D**, Schematic of the open field test. **E**, Repeated morphine administration leads to a reduction in the time spent in the center of an open field relative to mice treated with saline during adolescence, and saline during adulthood. Adolescent THC exposure blunted this reduction. One-way ANOVA p = 0.027; pairwise t-tests, vehicle/saline vs. vehicle/morphine 22.66 s vs. 12.55 s, p = 0.030; vehicle/saline vs. THC/morphine 22.66 s vs. 15.85 s, p = 0.077; vehicle/morphine vs. THC/morphine 12.55 s vs. 15.85 s, p = 0.59. **F**, Schematic of the marble burying task. **G**, Repeated morphine administration leads to an elevation in the number of marbles buried, indicative of anxiety-like behavior relative to mice treated with saline during adolescence, and saline during adulthood. Adolescent THC exposure prevented this morphine-induced anxiety. One-way ANOVA p = 0.0006; pairwise t-tests, vehicle/saline vs. vehicle/morphine 0.93 vs. 3.67, p = 0.0048; vehicle/saline vs. THC/morphine 0.93 vs. 0.50, p = 0.83; vehicle/morphine vs. THC/morphine 3.67 vs. 0.50, p = 0.006. **H**, Schematic of the von Frey task of mechanical hypersensitivity. **I**, Mice given saline vs. THC during adolescence showed no difference in baseline mechanical sensitivity. Vehicle/morphine vs. THC/morphine 0.85 g vs 0.86 g, p = 0.93. **J**, Repeated morphine administration leads to a reduction in the 50% mechanical threshold, indicative of pain hypersensitivity relative to mice treated with saline during adolescence, and saline during adulthood. Adolescent THC exposure blunted this morphine-induced pain hypersensitivity. One-way ANOVA p = 0.0032; pairwise t-tests, vehicle/saline vs. vehicle/morphine 1.09 vs. 0.42, p = 0.0027; vehicle/saline vs. THC/morphine 1.09 vs. 0.69, p = 0.052; vehicle/morphine vs. THC/morphine 0.42 vs. 0.69, p = 0.30.

**Figure 3:**
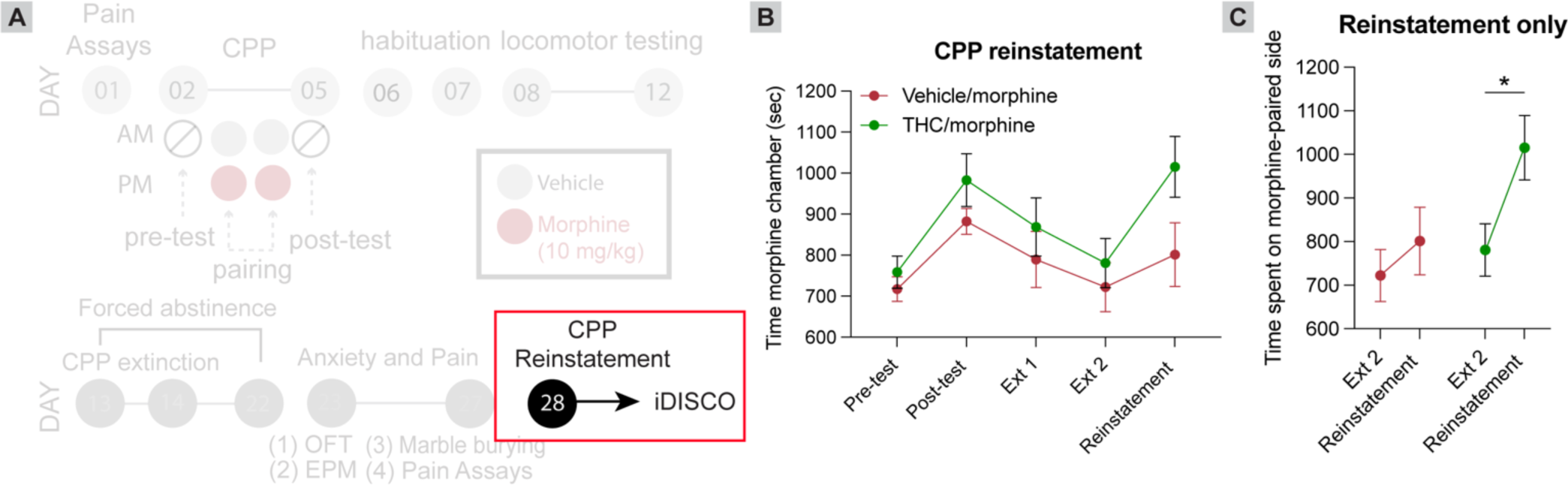
Mice treated with THC during adolescence show an elevation in drug-induced reinstatement of morphine-seeking behaviors later in life. **A**, Timeline of experiments. **B**, Time spent in the morphine-paired chamber during CPP pre-test, post-test, extinction, and reinstatement. Mice treated with THC or vehicle during adolescence showed a similar trajectory except for during reinstatement using a 5 mg/kg dose of morphine, at which time THC-treated mice showed an elevation in reinstatement behavior. **C**, Quantification of the time spent in the morphine-paired chamber in the reinstatement task relative to the final extinction day. Vehicle/morphine-treated mice 722 s vs 801.3 s, p = 0.43, n = 12 each; THC/morphine treated mice 780.7 s vs. 1015 s, p = 0.027, n = 8 each.

### Adolescent THC exposure impairs the development of anxiety-related behavior and pain hypersensitivity during morphine withdrawal

We next assessed whether mice treated with THC during adolescence were differentially impacted by withdrawal from repeated morphine administration. Following the five days of morphine given for locomotor testing and two days for CPP extinction, mice were left in their home cages for an additional 8 days, for a total abstinence period of ten days (Figure 2A). Following this ten-day forced abstinence, mice were tested for anxiety-like behavior using three tests: the elevated plus maze, open field test, and marble burying tests. The elevated plus maze tests how mice balance their tendency to explore against their preference for enclosed spaces (Figure 2B). A reduction in the time spent exploring the open arms is interpreted as anxiety-like behavior. Mice treated with repeated morphine injections spent less time in the open arms of the elevated plus maze relative to those treated with repeated saline injections. However, adolescent THC exposure blocked the reduction in time spent in the open arms induced by repeat morphine injection (One-way ANOVA p = 0.0005; pairwise t-tests, vehicle/saline vs. vehicle/morphine 47.73s vs. 17.84s, p = 0.01; vehicle/saline vs. THC/morphine 47.73s vs. 42.39s, p = 0.67; vehicle/morphine vs. THC/morphine 17.84s vs. 42.39s, p = 0.0018; Figure 2C). This suggests that adolescent THC exposure suppressed withdrawal-induced anxiety behaviors in response to adult opioid exposures. Similar results were observed using the open field test, where mice are both compelled to explore new environments, and avoid open areas (Figure 2D). Less time spent exploring the center of the arena is interpreted as anxiety-like behavior. We found that repeat morphine injections reduced time spent in the center of the open field, and THC history reduced this anxiety-like behavior (One-way ANOVA p = 0.027; pairwise t-tests, vehicle/saline vs. vehicle/morphine 22.66s vs. 12.55s, p = 0.030; vehicle/saline vs. THC/morphine 22.66s vs. 15.85s, p = 0.077; vehicle/morphine vs. THC/morphine 12.55s vs. 15.85s, p = 0.59; Figure 2E). Finally, in the marble burying test mice tend to bury marbles that are exposed in the bedding of a new cage (Figure 2F). In this assay, more marbles buried is interpreted to reflect increased anxiety-like behavior. We found that mice undergoing forced abstinence following repeat morphine injections buried more marbles after the 30-minute test, whereas adolescent THC administration reduced this number, consistent with a reduced anxiety-like behavior (One-way ANOVA p = 0.0006; pairwise t-tests, vehicle/saline vs. vehicle/morphine 0.93 vs. 3.67, p = 0.0048; vehicle/saline vs. THC/morphine 0.93 vs. 0.50, p = 0.83; vehicle/morphine vs. THC/morphine 3.67 vs. 0.50, p = 0.006; Figure 2G). Therefore, the results from each of these tests support the conclusion that adolescent THC exposure decreases the expression of anxiety-like behaviors in morphine-treated mice.

Next, we used a von Frey assay to examine the effects of adolescent THC exposure on mechanical hypersensitivity, both before and after morphine injection (Figure 2H). In addition to an increase in anxiety-like behavior, mice undergoing physiological withdrawal following repeated opioid administration show an increased pain sensitivity^42,43^. Mice treated with repeated THC doses during adolescence showed no difference in mechanical thresholds when tested one day prior to morphine CPP testing (Vehicle/morphine vs. THC/morphine 0.85g vs 0.86g, p = 0.93; Figure 2I). However, when tested again following repeat morphine exposure and forced abstinence, mice given repeated morphine administration showed an increased mechanical hypersensitivity relative to saline-treated controls, and adolescent THC treated reduced the extent of this hypersensitivity (One-way ANOVA p = 0.0032; pairwise t-tests, vehicle/saline vs. vehicle/morphine 1.09g vs. 0.42g, p = 0.0027; vehicle/saline vs. THC/morphine 1.09g vs. 0.69g, p = 0.052; vehicle/morphine vs. THC/morphine 0.42g vs. 0.69g, p = 0.30; Figure 2J). Together, these results indicate that adolescent THC exposure reduces the expression of morphine withdrawal-induced behaviors following repeated morphine administration during adulthood.

### Adolescent THC exposure enhances drug-induced reinstatement of morphine CPP

Following forced abstinence, we next assessed whether mice differentially reinstated CPP following a 5 mg/kg exposure of morphine in the CPP box. Drug exposure following abstinence can reinstate previously extinguished drug-seeking behaviors, and therefore can serve as a model of relapse in mice. Both groups of morphine-treated mice showed normal CPP (e.g., Figure 1D). Following CPP, mice then underwent extinction. To reduce confounding factors while examining reinstatement, mice were excluded that did not show CPP and proper extinction (N = 16 vehicle, 11 THC; Figure 3A-B). While mice treated with vehicle during adolescence did not significantly reinstate CPP following the priming 5 mg/kg morphine injection using this protocol, mice treated with THC during adolescence did significantly reinstate their CPP (Vehicle/morphine-treated mice 722s vs 801.3s, = p = 0.43; THC/morphine treated mice 780.7s vs. 1015s, p = 0.027; Figure 3C). This means that adolescent THC exposure drives reinstatement of drug-seeking behavior in response to a sub-threshold dose of morphine in mice given vehicle during adolescence.

### Whole-brain analysis shows differences in brain activity following drug-primed reinstatement in adolescent vehicle-vs. THC-treated mice

Given that adolescent THC-treated mice showed morphine-induced reinstatement whereas vehicle-treated controls did not, we next assessed what neural pathways may be responsible for this effect. To do this, we perfused the mice 60 minutes following the reinstatement test, cleared the brains using iDISCO+, and immunostained for the immediately early gene cFos to obtain a brain-wide map of neuronal activity (Figure 4A). We observed an overall elevation of neuronal activity in response to 5 mg/kg morphine in mice treated with THC during adolescence than vehicle, as evidenced by a higher number of cFos+ cells upon morphine exposure (Figure 4B). This increase was exhibited in most brain regions, with some of the highest-fold elevations occurring in basal ganglia regions including the globus pallidus externus (GPe), subthalamic nucleus (STN) and substantia nigra pars reticulata (SNr), as well as areas such as the substantia innominate/ventral pallidum, and several thalamic regions (Figure 4B-D). In fact, only four brain regions showed a reduction in cFos labeling in adolescent THC-treated mice, which included both the medial and lateral habenula.

**Figure 4:**
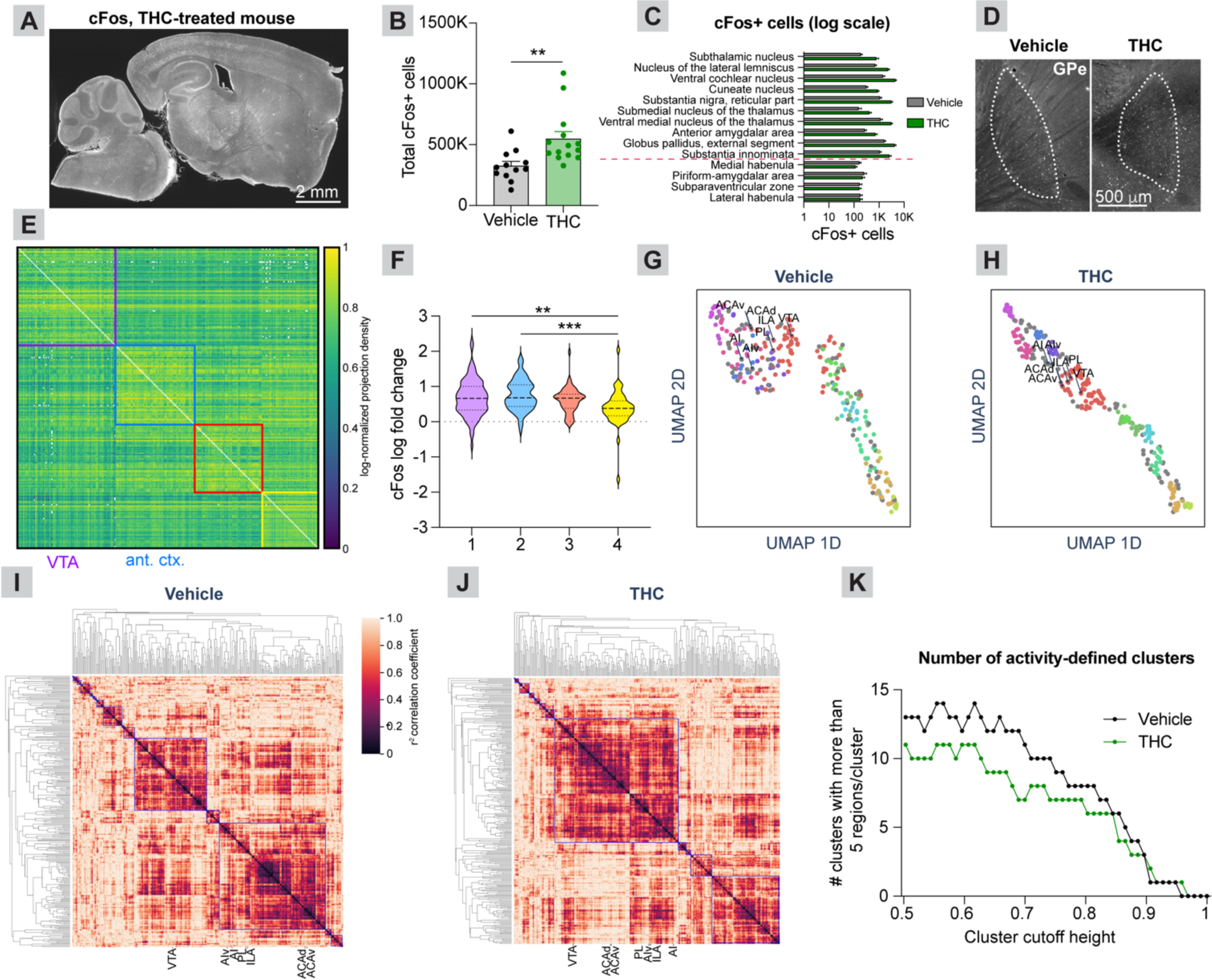
Brain-wide activity patterns in adolescent vehicle- and THC-treated mice following a reinstatement dose of morphine. **A**, Representative sagittal section of a cFos-stained brain of an animal treated with THC during adolescence and following a reinstatement dose of morphine. Scale, 2 mm. **B**, More cFos-expressing cells were detected in adolescent THC-treated mice than vehicle-treated mice following a dose of 5 mg/kg morphine (548,481 vs. 325,789 cells, p = 0.0051). **C**, The top ten brain regions showing the highest-fold elevation in cFos labeling in adolescent THC-treated relative to vehicle-treated mice and the four regions showing an elevation in vehicle-treated mice. Groups are separated by the dashed red line. Shown are brain regions with an average > 150 cFos+ cells in each brain region. **D**, Representative sagittal images of the GPe in adolescent vehicle-treated vs. THC-treated mice. Scale, 500 μm. **E**, Anatomical connectivity matrix of the mouse brain, broken into four communities based on the Louvain method for community detection **F**, Log fold-change in cFos labeling in each network community relative to one another (One-way ANOVA p = 0.0002; purple (1) vs. blue (2) 0.68 vs. 0.75, p = 0.76; purple (1) vs. red (3) 0.68 vs. 0.61, p = 0.82; purple (1) vs. yellow (4) 0.68 vs. 0.38, p = 0.0024; blue (2) vs. red (3) 0.75 vs. 0.61, p = 0.31; blue (2) vs. yellow (4) 0.68 vs. 0.38, p = 0.0002; red (3) vs. yellow (4) 0.61 vs. 0.38, p = 0.05. **G**, UMAP embedding of regions based on their correlated activity in adolescent vehicle-treated mice. Each point is a brain region. **H**, UMAP embedding of regions based on their correlated brain activity in adolescent THC-treated mice. The regions are overall more tightly clustered than in panel G, suggesting a greater level of correlated activity in adolescent THC-treated mice. **I**, Hierarchical clustering of regions based on the cFos correlation matrix in adolescent vehicle-treated mice. The VTA is within a separate activity-defined cluster from the frontal cortex regions of interest (prelimbic cortex, PL; infralimbic cortex, ILA; anterior insula cortex, AI; anterior insula cortex, ventral part, AIv; ACAd, anterior cingulate cortex, dorsal part; ACAv, anterior cingulate cortex, ventral part). **J**, Hierarchical clustering of regions based on the cFos correlation matrix in adolescent THC-treated mice. The VTA co-clusters with regions of the anterior cortex in these mice. **K**, The number of hierarchical clusters in adolescent vehicle-treated and THC-treated mice across a range of heights for cluster cut-offs in hierarchical trees. The THC-treated mice show reduced modularity across a range of cutoff values.

As the elevation in cFos labeling occurred globally in the mouse brain, we wanted to assess whether this elevation was larger for some pathways or whether the activation was uniform. To test this, we built a quantitative connectivity matrix of the mouse brain, using the Allen Mouse Brain Connectivity Atlas as a template^33^. Doing so revealed the existence of four major network communities within the brain, largely consisting of subcortical regions including the VTA (community 1, purple), cortical regions (community 2, blue), the hippocampus and related connections (community 3, red), and mid/hindbrain regions (community 4, yellow) (Figure 4E). We then assessed the relative increase in cFos labeling of regions in each community in adolescent THC-vs. vehicle-treated mice following a reinstatement opioid dose. We found a significant increase in the log fold-change cFos labeling in the purple and blue communities relative to the yellow community, suggesting that the subcortical and cortical modules both showed an overall larger increase in activity relative to mid/hindbrain regions in THC-treated mice (Figure 4F).

Following this analysis, we wanted to assess if these community-specific changes in cFos activation were accompanied by any changes in the functional relationships of different brain regions. To do so, we took the cFos labeling data across all regions for all mice and performed a dimensionality reduction analysis using Uniform Manifold Approximation and Projection (UMAP). UMAP divided the brain regions into approximately 10 activity-defined clusters in the adolescent THC-treated mice data. These clusters which mapped relatively consistently onto the adolescent vehicle-treated mice data (Figure 4G-H), with a larger distribution of brains from vehicle-treated compared to THC-treated mice. Interestingly, while frontal cortex regions, including the mPFC subregions infralimbic cortex (IL), prelimbic cortex (PL), anterior cingulate cortex (ACC), and anterior insula cortex (AI) all were in relatively close proximity in UMAP space in both adolescent vehicle-treated mice and THC-treated mice, they clustered much more tightly together in THC-treated group relative to the vehicle-treated group, indicating a higher level of correlated activity following a reinstatement opioid exposure (Figure 4G-H). This suggests that adolescent THC exposure may increase the functional connectivity between frontal cortex regions and the VTA. To assess functional modularity more quantitatively, we built a brain-wide functional correlation matrix for adolescent THC-treated and vehicle-treated mice based on their cFos labeling (Figure 4I-J). While in vehicle-treated mice the VTA and frontal cortex regions separated into different clusters, reflecting their separation into different networks (Figure 4I), in THC-treated mice the IL, PL, ACC, and AI all co-clustered with the VTA, indicating a higher level of correlated activity amongst these regions in adolescent THC-treated mice (Figure 4J). Last, we wanted to assess the overall modularity of brain activity in adolescent THC-vs. vehicle-treated mice. To do this, we determined the number of unique clusters across a range of parameters. Across a wide parameter range, adolescent THC-treated mice exhibited fewer activity-defined clusters than vehicle-treated mice, reflecting an overall reduced modularity in brains from adolescent THC-treated mice (Figure 4K). These findings overall show that the increased activation of cortical and subcortical pathways in adolescent THC-treated vs vehicle-treated mice after reinstatement is accompanied by a collapse in modularity of pathways in adolescent THC-treated mice, including a tighter clustering of the VTA with frontal cortex regions.

### Adolescent THC exposure changes connectivity to ventral tegmental area dopamine cells

VTA is a critical brain region for the development of a variety of drug-induced behavioral adaptations, including drug reward, sensitization, and withdrawal. To test whether adolescent THC exposure modified the input control to VTA dopamine (VTA^DA^) cells, we mapped inputs to these cells using the rabies virus (RABV) monosynaptic input mapping method. First, in animals treated with THC during adolescence, Cre-dependent adeno-associated viruses (AAVs) expressing TVA and the RABV glycoprotein, RABV-G, were injected into the VTA of DAT-Cre mice. Two weeks later, mice were injected with a glycoprotein-deleted, EnvA-pseudotyped, GFP-expressing RABV (Figure 5A). This virus infected TVA-expressing cells by nature of the EnvA-TVA interaction. Once in VTA^DA^ cells, this virus could spread to their inputs, as RABV-G was expressed in DAT-Cre cells. However, once in input cells, since RABV did not express its own glycoprotein, the virus could not spread further. Five days later, experiments were terminated, and GFP-positive cells were counted throughout the brain as we did previously^19,39^, and input connectivity was assessed. RABV-labeled input cells were observed throughout the brain, as in previous studies (Figure 5B-C)^19,39^. Overall, we observed several quantitative differences, shown in Figure 5D (Two-way ANOVA, interaction p = 0.0062). While several brain regions showed a visible difference in connectivity, for example elevated connectivity from cortical regions comprising the anterior cortex (infralimbic cortex, prelimbic cortex, anterior insula cortex, anterior cingulate cortex, and the anterior motor cortex^19,38,39,44^), parabrachial nucleus (PBN) and deep cerebellar nuclei (DCN), only the anterior cortex was significantly different following correction for multiple comparisons (corrected p-value p = 0.029).

**Figure 5:**
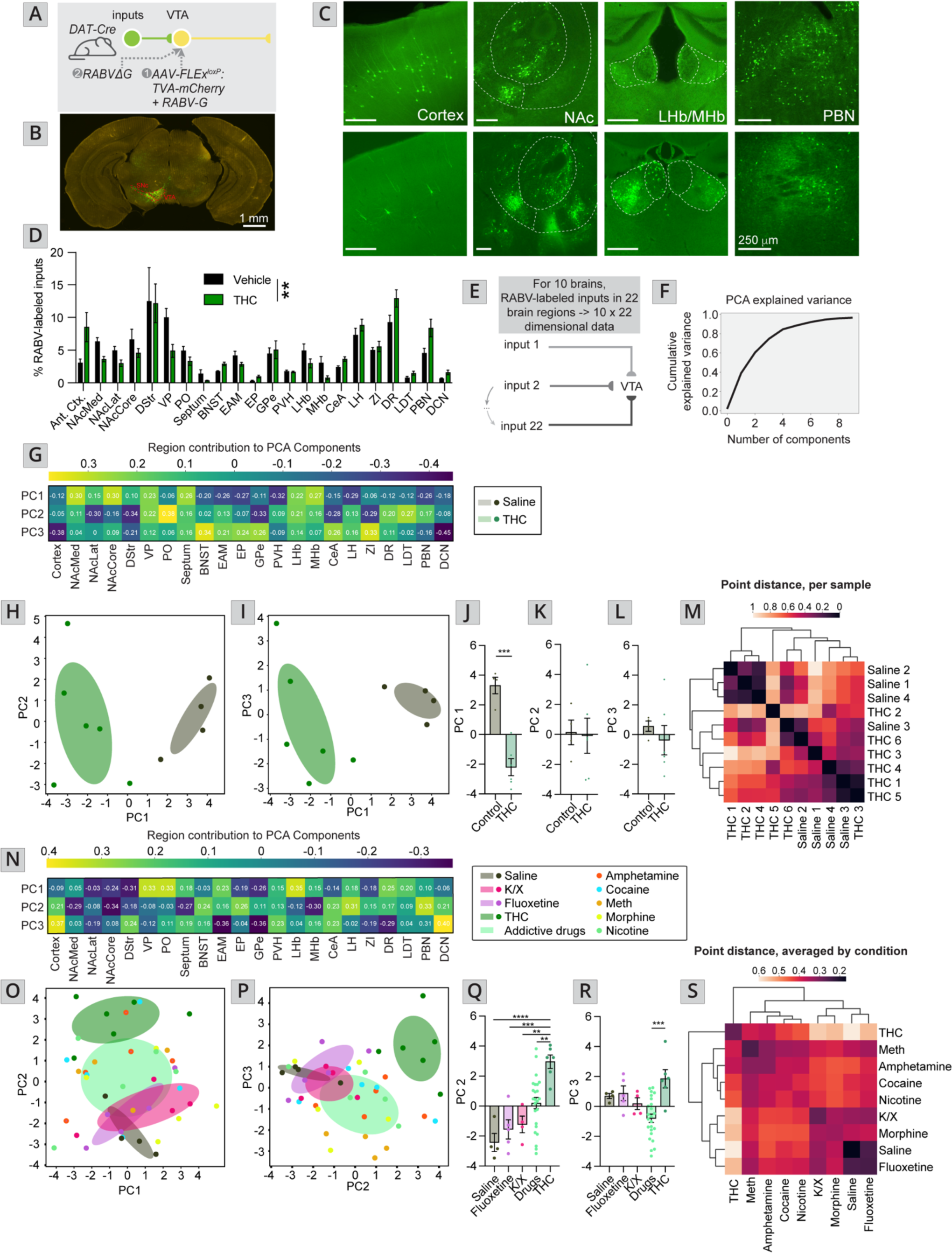
THC-treated mice show a different brain-wide pattern relative to animals treated with psychostimulants, opioids, or nicotine. **A**, Schematic of experiments. **B**, Representative image of the ventral midbrain of a DAT-Cre mouse showing starter cells in the VTA and medial SNc. Scale, 1 mm. **C**, Representative images of input cell populations in several brain sites, including the anterior cortex, NAcMed, NAcCore, NAcLat, MHb, LHb, and PBN. Scale, 250 μm. **D**, Bar graph plot showing the percentage of RABV-labeled inputs in control vs. THC-treated mice. Two-way ANOVA, interaction p = 0.0062. Only the anterior cortex was significantly different following correction for multiple comparisons (corrected p-value p = 0.029. **E**, Schematic and explanation for how PCA analysis was carried out on the RABV tracing dataset. **F**, Percent of cumulative variance explained by each principal component. **G**, Heatmap of the contributions of each brain region, or feature, in the data to PCs 1 through 3. **H**, Plot of PC1 and PC2 from brains of control and THC-treated mice. **I**, Plot of PC1 and PC3 from brains of control and THC-treated mice. **J-L**, Comparison of total PC values for each brain along **J** PC 1, **K** PC 2, and **L** PC 3. **J**, Control 3.3 vs. THC -2.2, p = 0.0002. **K**, Control 0.13 vs. THC -0.09, p = 0.89. **L**, Control 0.57 vs. THC -0.38, p = 0.48. **M**, To further quantify relationships between brains and conditions, the Euclidean distance between each brain from control and THC-treated mice in the PC1 versus PC2 (or PC1 vs. PC3) coordinate space was calculated. These distances were then plotted in a heatmap with brains being organized by similarity assessed by hierarchical clustering. **N**, Heatmap of the contributions of each brain region, or feature, in the data to PCs 1 through 3 in the data presented in panels N-Q. **O**, Plot of PC1 and PC2 from brains of control and drug-treated mice. **P**, Plot of PC1 and PC3 from brains of control and drug-treated mice. **Q-R**, Comparison of total PC values for each brain along **Q** PC 2, and **R** PC 3. For PC 1, One-way ANOVA p = 0.83. **Q**, One-way ANOVA p < 0.0001; pairwise t-tests and asterisks presented in the figure include THC comparisons only, although statistics are based on comparisons between all groups. Saline vs. THC -2.43 vs. 2.96, p < 0.0001; Fluoxetine vs. THC -1.55 vs. 2.96, p = 0.0004; K/X vs. THC -1.23 vs. 2.96 p = 0.0024; Drugs vs. THC 0.19 vs. 2.96, p = 0.0051. **R**, One-way ANOVA p = 0.0004; Saline vs. THC 0.70 vs. 1.85, p = 0.63; Fluoxetine vs. THC 0.88 vs. 1.85, p = 0.71; K/X vs. THC 0.19 vs. 1.85 p = 0.27; Drugs vs. THC -0.79 vs. 1.85, p = 0.0005. **S**, The Euclidean distance between each psychoactive drug-treated and control brain in the PC1 versus PC2 coordinate space was calculated, plotted in a heatmap, and brains were organized by similarity and assessed by hierarchical clustering.

This traditional approach to assessing differences in connectivity is statistically conservative, and only assesses potential differences in connectivity, one input at a time. To reveal further patterns in the data, we have recently employed a dimensionality reduction approach that enables us to detect larger-scale differences in input patterns between brains of mice undergoing a variety of treatments^39,45^. We therefore first used this approach to assess which inputs best differentiated THC-treated vs. control mice using principal component analysis (PCA). We found that 3 PCs were sufficient to explain approximately 75% of the variance in the data, and thus we focused only on these 3 PCs (Figure 5E-F). We found that PC1 completely separated THC-treated and control mice, whereas PC2 and PC3 did not (Figure 5G-I). Control mice had high PC1 values, which were driven predominately by strong contributions from nucleus accumbens medial shell (NAcMed), core (NAcCore), ventral pallidum (VP), septum, lateral habenula (LHb) and medial habenula (MHb), while THC-treated mice had low PC1 values, which were driven by the bed nucleus of the stria terminalis (BNST), extended amygdala area (EAM), entopeduncular nucleus (EP), paraventricular nucleus of the hypothalamus (PVH), lateral hypothalamus (LH), and PBN (Figure 5G). To quantify these differences, we plotted each brain’s location on each PC, and performed t-tests. Differences between THC-treated and control groups were strongly significant along PC1, and not different along PCs 2 and 3, as expected (Control 3.3 vs. THC -2.2, p = 0.0002). To further quantify relationships between brains and conditions, the Euclidean distance between each brain in the PC1 versus PC2 coordinate space was calculated, and the relationships plotted as a correlogram. THC-treated and control brains separated largely by condition, as expected (Figure 5M).

Last, we assessed how VTA^DA^ connectivity of THC-treated mice compares to mice treated with one of several drugs, including a single dose of the psychoactive but non-addictive fluoxetine, two doses of the anesthetic mixture of 100 mg/kg ketamine and 10 mg/kg xylazine, spaced 2 weeks apart, or one dose of the addictive cocaine, amphetamine, methamphetamine, morphine, or nicotine, together grouped for simplicity as “drugs of abuse”. Using a similar PCA-based assessment, we observed that mice treated with a drug of abuse separated from the other conditions along PC2, but not PC1 or PC3 (Figure 5N-P). Interestingly, THC-treated mice had even higher values of PC2 than mice treated with one of the drugs of abuse and were thus located further along PC2 from controls (Figure 5O-P). Positive values of PC2 were largely driven by strong contributions from the anterior cortex, BNST, EP, CeA, LH, PBN, and DCN, notably many of the same regions that differentiated brains of THC-treated from controls (Figure 5G, N). While PC3 did not strongly differentiate most of the groups, it did differentiate THC-treated mice from those treated with a drug of abuse (Figure 5R). THC-treated mice had higher PC3 values, which were driven by strong contributions from the anterior cortex, dorsal striatum (DStr), PVH, CeA, PBN, and DCN (Figure 5N). The locations of each brain in PC space were plotted on bar graphs, and differences were again assessed (PC2; Saline vs. THC -2.43 vs. 2.96, p < 0.0001; Fluoxetine vs. THC -1.55 vs. 2.96, p = 0.0004; K/X vs. THC -1.23 vs. 2.96 p = 0.0024; Drugs vs. THC 0.19 vs. 2.96, p = 0.0051. PC3; One-way ANOVA p = 0.0004; Saline vs. THC 0.70 vs. 1.85, p = 0.63; Fluoxetine vs. THC 0.88 vs. 1.85, p = 0.71; K/X vs. THC 0.19 vs. 1.85 p = 0.27; Drugs vs. THC -0.79 vs. 1.85, p = 0.0005; Figure 5Q-R). The distinct connectivity patterns were also apparent following our Euclidean distance analysis in PC1 and PC2 space, which indicated that the connectivity patterns from brains from THC-treated mice segregate completely from those of mice treated with other psychoactive drugs (Figure 5S). Together, these results indicate that adolescent THC treatment induces similar but distinct input changes onto VTA^DA^ neurons as previous exposure to a single exposure to a “hard drug”, principally an increase in connectivity from the anterior cortex.

## DISCUSSION

Here, we tested the effects of adolescent THC exposure on later-life morphine-induced behaviors, brain-wide activity following morphine reinstatement, and midbrain dopamine cell synaptic connectivity. We found that adolescent THC exposure blunts morphine-induced locomotion and withdrawal anxiety/pain hypersensitivity and enhances morphine-induced reinstatement following a forced abstinence period. This broad range of effects is consistent with findings from studies that have explored the influence of THC on morphine-induced behavioral changes^14,25–27,46^. Here, we extend these previous studies by linking brain-wide activity patterns following a reinstatement dose of morphine to morphine-induced reinstatement behavior and show that adolescent THC exposure creates long-lasting changes in input connectivity to VTA^DA^ cells. Our study thus adds to a rich literature on the subject by implicating several discrete circuits as potential drivers of the long-lasting effects of adolescent THC exposure on behavioral and brain responses to opioids later in life.

It is perhaps surprising that THC exposure blunts morphine withdrawal-induced anxiety across several behavioral assays, as well as pain sensitivity. Interestingly, the behavioral profile is consistent with the behavioral responses observed following exposure of drugs of abuse during adolescence^47–51^, where adolescent mice experience potentiated drug reward with only mild withdrawal symptoms relative to adult mice receiving equivalent drug exposures. This effect could be mediated by an occlusion of opioid-induced epigenetic changes by those elicited following adolescent THC exposure. Such an effect was previously observed following juvenile exposure to oxycodone; in this case, oxycodone induced long-lasting epigenetic changes, for example an elevation of the repressive histone modification H3K27me3 on genes that control dopamine transmission in the VTA^52^. Therefore, one hypothesis is that adolescent THC exposure may result in a similar imbalance of reward/aversion as is seen in responses to drugs during adolescence by occluding compensatory responses to drug exposure that typically occur in adulthood, but not adolescence. This could result, in, for example, withdrawal from repeated opioid injection failing to induce a compensatory induction in anxiety-related behaviors and pain hypersensitivity. In this case, reversal of these epigenetic marks may restore mice to basal levels of susceptibility. Future studies are required to identify the potential cellular loci of these effects, as well as what the epigenetic consequences of early life THC exposure may be.

In addition, we found that adolescent THC administration induces higher levels of drug-induced reinstatement, consistent with previous observations assessing drug- or stress-induced reinstatement^8,9,26^. Here, we extend these observations by linking these results to brain-wide cFos activation in adolescent vehicle-vs. THC-treated mice. Overall, we found that adolescent THC-treated mice had higher numbers of cFos+ cells in response to a 5 mg/kg reinstatement dose of morphine than adolescent vehicle-treated controls. In addition, we found that adolescent THC-treated mice have higher levels of correlated activity in mPFC and AI regions with the VTA than adolescent vehicle-treated mice, indicating enhanced functional connectivity between these regions (Figure 4G-H). We previously showed that activation of these cortical inputs to the VTA promotes reinforcement through increasing dopamine release in the NAc^38^. Therefore, the elevated functional connectivity that we observe between these regions in adolescent THC-treated mice may promote greater levels of dopamine release in the NAc than in adolescent vehicle-treated mice and is one mechanism by which adolescent THC-treated mice may more strongly reinstate morphine CPP following a priming dose of morphine.

However, one limitation of cFos-based analyses is that they do not consider anatomical connections. Viral methods such as the monosynaptic RABV circuit mapping technique can specifically label synaptically-connected cells^53,54^. Furthermore, we recently showed that the extent of input labeling is dependent on the activity of input connections^19,20,55^. Therefore, our RABV mapping experiments have the unique ability to explore the changes in connectivity to VTA^DA^ cells following repeated THC exposure, without a preconceived bias as to which inputs may be altered. We have previously used this assay to define how drugs of abuse change inputs to VTA^DA^ cells, enabling us to identify several pathway-specific changes that have important consequences for the development of drug-induced behavioral adaptations^39^. Here, we compare connectivity of VTA^DA^ cells in brains from mice receiving our adolescent THC protocol to those receiving a single drug of abuse. We found that adolescent THC exposure induced numerous long-lasting input changes onto VTA^DA^ cells that partially, but not completely, overlapped with those induced by “hard drugs”. The most prominent difference between mice given adolescent THC exposure vs. a drug of abuse is that THC-treated mice show a pronounced increase in inputs from the anterior cortex (Figure 5D). Heightened activation of this connection is consistent with our cFos results and would be expected to be rewarding through elevating activity in VTA^DA^ cells. This hypothesis is consistent with the hyper-dopaminergic state of THC-treated animals previously reported^15^, and the more robust response in these animals to a reinstatement dose of morphine. In addition, we observed an overall reduced modularity in adolescent THC-mice relative to adolescent vehicle-treated controls (Figure 4K). A reduction in brain modularity is seen following use and withdrawal from drugs of abuse^56–61^ as well as brain disorders such as traumatic brain injury and Alzheimer’s-related dementia^62–67^. How this reduced modularity may influence behavioral responses to drugs, however, remains largely unknown.

In our study, we also tested the hypothesis that adolescent THC exposure alters connectivity onto VTA dopamine (VTA^DA^) cells. This hypothesis was supported by the observation that rats treated with THC during adolescence develop a hyper-dopaminergic state^15^; one potential mechanism that could mediate this effect would be a change in connectivity to facilitate an increase in DA release in downstream brain regions. Here, we found an elevation in inputs from the anterior cortex to VTA^DA^ cells, which we previously showed promotes reinstatement by facilitating DA release in the NAc. Notably, we previously have shown that a single injection of a variety of drugs of abuse can cause long-lasting changes in input connectivity onto VTA^DA^ cells^19,20,55^, with this signature being relatively similar regardless of the drug administered^19,39^, suggesting that these drugs induce similar changes in connectivity. The anterior cortex, particularly the mPFC, is thought to play important roles in the later stages of substance use disorder^68–71^. The reduction in connectivity from the anterior following a single dose, and increase in connectivity following repeated doses, is thus consistent with that hypothesis of altered mPFC engagement during the addiction cycle. One additional point of note is that we previously showed that drugs of abuse increase connectivity from the globus pallidus externus (GPe) to the ventral midbrain, and that these changes are important for drug-induced behavioral adaptations including reward and sensitization^19,20^. However, THC treatment did not induce a notable change in GPe inputs onto VTA^DA^ cells (Figure 5D). Thus, while THC triggers several long-lasting changes in connectivity to VTA^DA^ cells, these appear to be distinct from those initiated by psychostimulants, nicotine, or morphine. However, the conclusions that we can draw from these experiments are limited by differences in experimental design. For example, for the drug-treated mice, RABV was injected one day following injection of the drug, whereas in THC-treated mice, they were injected approximately 35 days following completion of a 2-week daily injection protocol. Thus, while in the former study design drug-induced changes do not need to persist for longer than a few days to yield the observed changes, in THC-treated animals the effects observed had to persist for longer than 35 days. In addition, THC was given fourteen times, once per day, whereas all other drugs were only given once in total. Therefore, the effects observed with THC may result from repeated injections. However, we performed the experiments presented here because our behavioral assessments were done 35 days following completion of THC administration, and we wanted the RABV experiments to follow the same timeline. Further experiments will be needed to resolve these questions.

## Acknowledgements

We would like to thank Dr. Daniele Piomelli and the Impact of Cannabinoids Across the Lifespan team at the University of California, Irvine for providing seed funding for this work. This work was additionally funded by the NIH (R00 DA041445, DP2 AG067666, R01NS130044, R01DA056599, 1R01DA054374), Tobacco Related Disease Research Program (T31KT1437, T31P1426), One Mind (OM-5596678), Brightfocus Foundation (A2022031S), and the Brain and Behavior Research Foundation (NARSAD 26845) to KTB, T32 GM136624 to KB and PD, NSF GRFP DGE-1839285 to PD, T32DA050558 to EH, and R01DA055849, U01DA053826, R01MH132680, and P50DA044118 to SVM.

## REFERENCES

1. National Center for Drug Abuse Statistics. Drug Abuse Statistics. (2024).

2. Hasin, D. S. et al. Prevalence of marijuana use disorders in the United States between 2001-2002 and 2012-2013. JAMA Psychiatry vol. 72 1235–1242 Preprint at 10.1001/jamapsychiatry.2015.1858 (2015).

3. Ferland, J. M. N. & Hurd, Y. L. Deconstructing the neurobiology of cannabis use disorder. Nat Neurosci 23, 600–610 (2020).

4. Ruiz, C. M. et al. Pharmacokinetic, behavioral, and brain activity effects of ?9-tetrahydrocannabinol in adolescent male and female rats. Neuropsychopharmacology 46, 959–969 (2021).

5. Chadwick, B., Miller, M. L. & Hurd, Y. L. Cannabis Use during Adolescent Development: Susceptibility to Psychiatric Illness. Front Psychiatry 4, (2013).

6. Rubino, T., Zamberletti, E. & Parolaro, D. Adolescent exposure to cannabis as a risk factor for psychiatric disorders. Journal of Psychopharmacology vol. 26 177–188 Preprint at 10.1177/0269881111405362 (2012).

7. Panlilio, L. V., Zanettini, C., Barnes, C., Solinas, M. & Goldberg, S. R. Prior exposure to THC increases the addictive effects of nicotine in rats. Neuropsychopharmacology 38, 1198–1208 (2013).

8. Stopponi, S. et al. Chronic THC during adolescence increases the vulnerability to stress-induced relapse to heroin seeking in adult rats. European Neuropsychopharmacology 24, 1037–1045 (2014).

9. Lecca, D. et al. Adolescent cannabis exposure increases heroin reinforcement in rats genetically vulnerable to addiction. Neuropharmacology 166, (2020).

10. Ellgren, M., Spano, S. M. & Hurd, Y. L. Adolescent cannabis exposure alters opiate intake and opioid limbic neuronal populations in adult rats. Neuropsychopharmacology 32, 607–615 (2007).

11. Panlilio, L. V., Solinas, M., Matthews, S. A. & Goldberg, S. R. Previous exposure to THC alters the reinforcing efficacy and anxiety-related effects of cocaine in rats. Neuropsychopharmacology 32, 646–657 (2007).

12. Friedman, A. L., Meurice, C. & Jutkiewicz, E. M. Effects of adolescent Δ9-tetrahydrocannabinol exposure on the behavioral effects of cocaine in adult Sprague–Dawley rats. Exp Clin Psychopharmacol 27, 326–337 (2019).

13. Lüscher, C. & Malenka, R. C. Drug-Evoked Synaptic Plasticity in Addiction: From Molecular Changes to Circuit Remodeling. Neuron 69, 650–663 (2011).

14. Rubino, T. et al. The Depressive Phenotype Induced in Adult Female Rats by Adolescent Exposure to THC is Associated with Cognitive Impairment and Altered Neuroplasticity in the Prefrontal Cortex. Neurotox Res 15, 291–302 (2009).

15. Renard, J. et al. Adolescent Cannabinoid Exposure Induces a Persistent Sub-Cortical Hyper-Dopaminergic State and Associated Molecular Adaptations in the Prefrontal Cortex. Cereb Cortex 27, 1297–1310 (2017).

16. Renard, J. et al. Adolescent THC Exposure Causes Enduring Prefrontal Cortical Disruption of GABAergic Inhibition and Dysregulation of Sub-Cortical Dopamine Function /631/378/2571/631/378/1689/1799 /9 /9/30 /82 /82/1 article. Sci Rep 7, (2017).

17. Creed, M. et al. Cocaine exposure enhances the activity of ventral tegmental area dopamine neurons via calcium-impermeable NMDARs. Journal of Neuroscience 36, 10759–10768 (2016).

18. Bocklisch, C. et al. Cocaine Disinhibits Dopamine Neurons by Potentiation of GABA Transmission in the Ventral Tegmental Area. Science (1979) 341, 1521–1525 (2013).

19. Beier, K. T. et al. Rabies screen reveals GPe control of cocaine-triggered plasticity. Nature 549, 345–350 (2017).

20. Tian, G. et al. Molecular and circuit determinants in the globus pallidus mediating control of cocaine-induced behavioral plasticity. Neuron Sneak Peek (2023).

21. Kandel, D. B. Marijuana Users in Young Adulthood. Arch Gen Psychiatry 41, 200–209 (1984).

22. Clayton, R. & Voss, H. Young men and drugs in Manhattan: a causal analysis. NIDA Res. Monogr. 39, 1–187 (1981).

23. O’Donnell, J. & Clayton, R. The stepping-stone hypothesis--marijuana, heroin, and causality. Chem Depend. 4, 229–241 (1982).

24. Tanda, G., Pontieri, F. E. & Chiara, G. Di. Cannabinoid and Heroin Activation of Mesolimbic Dopamine Transmission by a Common 1 Opioid Receptor Mechanism. https://www.science.org.

25. Cadoni, C., Valentini, V. & Di Chiara, G. Behavioral sensitization to Δ9-tetrahydrocannabinol and cross-sensitization with morphine: Differential changes in accumbal shell and core dopamine transmission. J Neurochem 106, 1586–1593 (2008).

26. Ellgren, M., Spano, S. M. & Hurd, Y. L. Adolescent cannabis exposure alters opiate intake and opioid limbic neuronal populations in adult rats. Neuropsychopharmacology 32, 607–615 (2007).

27. Tomasiewicz, H. C. et al. Proenkephalin mediates the enduring effects of adolescent cannabis exposure associated with adult opiate vulnerability. Biol Psychiatry 72, 803–810 (2012).

28. Cicero, T. J., Ellis, M. S., Surratt, H. L. & Kurtz, S. P. The changing face of heroin use in the United States a retrospective analysis of the past 50 years. JAMA Psychiatry 71, 821–826 (2014).

29. Muhuri, P., Gfroerer, J. & Davies, M. Substance Abuse and Mental Health Services Administration. Associations of Nonmedical Pain Reliever Use and Initiation of Heroin Use in the United States. (2013).

30. Chaplan, S. R., Bach, F. W., Pogrel, J. W., Chung, J. M. & Yaksh, T. L. Quantitative assessment of tactile allodynia in the rat paw. J Neurosci Methods 53, 55–63 (1994).

31. Renier, N. et al. IDISCO: A simple, rapid method to immunolabel large tissue samples for volume imaging. Cell 159, 896–910 (2014).

32. Renier, N. et al. Mapping of Brain Activity by Automated Volume Analysis of Immediate Early Genes. Cell 165, 1789–1802 (2016).

33. Oh, S. W. et al. A mesoscale connectome of the mouse brain. Nature 508, 207–214 (2014).

34. Muller, B., Richards, A. J., Jin, B. & Lu, X. GOGrapher: A Python library for GO graph representation and analysis. BMC Res Notes 2, (2009).

35. McInnes, L., Healy, J. & Melville, J. UMAP: Uniform Manifold Approximation and Projection for Dimension Reduction. ArXiv >1802.03426, 1–51 (2018).

36. McInnes, L., Healy, J. & Astels, S. hdbscan: Hierarchical density based clustering. The Journal of Open Source Software 2, 205 (2017).

37. Waskom, M. Seaborn: Statistical Data Visualization. J Open Source Softw 6, 3021 (2021).

38. Beier, K. T. et al. Circuit Architecture of VTA Dopamine Neurons Revealed by Systematic Input-Output Mapping. Cell 162, 622–634 (2015).

39. Bartas, K. et al. Drug-induced changes in connectivity to midbrain dopamine cells revealed by rabies monosynaptic tracing. Elife 13, RP93664 (2024).

40. Cooper, Z. D. & Haney, M. Comparison of subjective, pharmacokinetic, and physiological effects of marijuana smoked as joints and blunts. Drug Alcohol Depend 103, 107–113 (2009).

41. Huestis, M. A., Henningfield, J. E. & Cone, E. J. Blood Cannabinoids. I. Absorption of THC and Formation of 11-OH-THC and THCCOOH During and After Smoking Marijuana*. J Anal Toxicol 16, 276–282 (1992).

42. Koob, G. F. Neurobiology of Opioid Addiction: Opponent Process, Hyperkatifeia, and Negative Reinforcement. Biol Psychiatry 87, 44–53 (2020).

43. Koob, G. F. Drug addiction: Hyperkatifeia/negative reinforcement as a framework for medications development. Pharmacol Rev 73, 163–201 (2021).

44. Beier, K. T. et al. Topological Organization of Ventral Tegmental Area Connectivity Revealed by Viral-Genetic Dissection of Input-Output Relations. Cell Rep 26, 159–167.e6 (2019).

45. Derdeyn, P., Hui, M., Macchia, D. & Beier, K. Uncovering the Connectivity Logic of the Ventral Tegmental Area. Front Neural Circuits 15, 799688 (2022).

46. Cadoni, C., Pisanu, A., Solinas, M., Acquas, E. & Di Chiara, G. Behavioural sensitization after repeated exposure to Δ9-tetrahydrocannabinol and cross-sensitization with morphine. Psychopharmacology (Berl) 158, 259–266 (2001).

47. Levin, E. D., Rezvani, A. H., Montoya, D., Rose, J. E. & Scott Swartzwelder, H. Adolescent-onset nicotine self-administration modeled in female rats. Psychopharmacology (Berl) 169, 141–149 (2003).

48. Faraday, M. M., Elliott, B. M. & Grunberg, N. E. Adult vs. Adolescent Rats Differ in Biobehavioral Responses to Chronic Nicotine Administration. www.elsevier.com/locate/pharmbiochembeh.

49. Vastola, B. J., Douglas, L. A., Varlinskaya, E. I. & Spear, L. P. Nicotine-Induced Conditioned Place Preference in Adolescent and Adult Rats.

50. Faraday MM, Elliott BM, Phillips JM & Grunberg NE. Adolescent and adult male rats differ in sensitivity to nicotine’s activity effects. Pharmacol Biochem Behav. 74, 917–931 (2003).

51. O’Dell, L. E., Bruijnzeel, A. W., Ghozland, S., Markou, A. & Koob, G. F. Nicotine withdrawal in adolescent and adult rats. in Annals of the New York Academy of Sciences vol. 1021 167–174 (New York Academy of Sciences, 2004).

52. Carpenter, M. D., Manners, M. T., Heller, E. A. & Blendy, J. A. Adolescent oxycodone exposure inhibits withdrawal-induced expression of genes associated with the dopamine transmission. Addiction Biology 26, (2021).

53. Wickersham, I. R. et al. Monosynaptic Restriction of Transsynaptic Tracing from Single, Genetically Targeted Neurons. Neuron 53, 639–647 (2007).

54. Wall, N. R., Wickersham, I. R., Cetin, A., De La Parra, M. & Callaway, E. M. Monosynaptic circuit tracing in vivo through Cre-dependent targeting and complementation of modified rabies virus. Proc Natl Acad Sci U S A 107, 21848–21853 (2010).

55. Tian, G. et al. An extended Amygdala-midbrain circuit controlling cocaine withdrawal-induced anxiety and reinstatement. Cell Rep 39, 1–16 (2022).

56. Kimbrough, A. et al. Brain-wide functional architecture remodeling by alcohol dependence and abstinence. PNAS 117, 2149–2159 (2020).

57. Orsini, C. A., Colon-Perez, L. M., Heshmati, S. C., Setlow, B. & Febo, M. Functional connectivity of chronic cocaine use reveals progressive neuroadaptations in neocortical, striatal, and limbic networks. eNeuro 5, (2018).

58. Konova, A. B., Moeller, S. J., Tomasi, D. & Goldstein, R. Z. Effects of chronic and acute stimulants on brain functional connectivity hubs. Brain Res 1628, 147–156 (2015).

59. Liang, X. et al. Interactions between the salience and default-mode networks are disrupted in cocaine addiction. Journal of Neuroscience 35, 8081–8090 (2015).

60. Tomasi, D. et al. Disrupted functional connectivity with dopaminergic midbrain in cocaine abusers. PLoS One 5, (2010).

61. Konova, A. B., Moeller, S. J., Tomasi, D., Volkow, N. D. & Goldstein, R. Z. Effects of methylphenidate on resting-state functional connectivity of the mesocorticolimbic dopamine pathways in cocaine addiction. JAMA Psychiatry 70, 857–868 (2013).

62. Bertolero, M. A., Yeo, B. T. T., Bassett, D. S. & D’Esposito, M. A mechanistic model of connector hubs, modularity and cognition. Nat Hum Behav 2, 765–777 (2018).

63. Gallen, C. L. et al. Modular brain network organization predicts response to cognitive training in older adults. PLoS One 11, (2016).

64. Brier, M. R. et al. Functional connectivity and graph theory in preclinical Alzheimer’s disease. Neurobiol Aging 35, 757–768 (2014).

65. De Haan, W. et al. Disrupted modular brain dynamics reflect cognitive dysfunction in Alzheimer’s disease. Neuroimage 59, 3085–3093 (2012).

66. Arnemann, K. L., et al. Functional Brain Network Modularity Predicts Response to Cognitive Training after Brain Injury. https://www.neurology.org (2015).

67. Sporns, O. & Betzel, R. F. Modular brain networks. Annu Rev Psychol 67, 613–640 (2016).

68. Koob, G. F. & Volkow, N. D. Neurocircuitry of addiction. Neuropsychopharmacology 35, 217–238 (2010).

69. Goldstein, R. Z. & Volkow, N. D. Dysfunction of the prefrontal cortex in addiction: Neuroimaging findings and clinical implications. Nat Rev Neurosci 12, 652–669 (2011).

70. Volkow, N. D., Michaelides, M. & Baler, R. The neuroscience of drug reward and addiction. Physiol Rev 99, 2115–2140 (2019).

71. Miller, E. K. & Cohen, J. D. AN INTEGRATIVE THEORY OF PREFRONTAL CORTEX FUNCTION. Annu. Rev. Neurosci 24, 167–202 (2001).

